# SARS-CoV-2 Delta Variant Remains Viable in Environmental Biofilms found in Meat Packaging Plants

**DOI:** 10.1101/2023.06.15.545172

**Authors:** Austin B. Featherstone, Arnold J. T. M. Mathijssen, Amanda Brown, Sapna Chitlapilly Dass

## Abstract

Severe acute respiratory syndrome coronavirus 2 (SARS-CoV-2) is a coronavirus that directly infects human airway epithelial cells and caused the COVID-19 pandemic. At the start of the pandemic in 2020, meat-packaging plants saw a surge in SARS-CoV-2 cases, which forced many to temporarily close. To determine why SARS-CoV-2 appears to thrive specifically well in meat packaging plants, we used SARS-CoV-2 Delta variant and meat packaging plant drain samples to develop mixed-species biofilms on materials commonly found within meat packaging plants (stainless steel (SS), PVC, and ceramic tile). Our data provides evidence that SARS-CoV-2 Delta variant remained viable on all the surfaces tested with and without an environmental biofilm. We observed that SARS-CoV-2 Delta variant was able to remain infectious with each of the environmental biofilms, however, we detected a significant reduction in viability post-exposure to Plant B biofilm on SS, PVC, and on ceramic tile chips, and to Plant C biofilm on SS and PVC chips. The numbers of viable SARS-CoV-2 Delta viral particles was 1.81 – 4.57-fold high than the viral inoculum incubated with the Plant B and Plant C environmental biofilm on SS, and PVC chips. We did not detect a significant difference in viability when SARS-CoV-2 Delta variant was incubated with the biofilm obtained from Plant A on any of the materials tested and SARS-CoV-2 Delta variant had higher plaque numbers when inoculated with Plant C biofilm on tile chips, with a 2.75-fold difference compared to SARS-CoV-2 Delta variant on tile chips by itself. In addition, we detected an increase in the biofilm biovolume in response to SARS-CoV-2 Delta variant which is also a concern for food safety due to the potential for foodborne pathogens to respond likewise when they come into contact with the virus. These results indicate a complex virus-environmental biofilm interaction which correlates to the different bacteria found in each biofilm. Our results also indicate that there is the potential for biofilms to protect SARS-CoV-2 from disinfecting agents and remaining prevalent in meat packaging plants. With the highly infectious nature of some SARS-CoV-2 variants such as Delta, and more so with the Omicron variant, even a minimal amount of virus could have serious health implications for the spread and reoccurrence of SARS-CoV-2 outbreaks in meat packaging plants.

## Introduction

Severe acute respiratory syndrome coronavirus 2 (SARS-CoV-2) belongs to the genus β coronaviruses. In 2019, a new strain of coronavirus (SARS-CoV-2) was discovered to directly infect humans without an animal reservoir and cause a severe respiratory disease in humans called Coronavirus Disease-2019 (COVID-19) [1–3]. The first SARS-CoV-2 wild-type (WT) cases recorded in the United States were found in Washington and in Illinois in January 2020 [4,5]. After the initial WT cases of SARS-CoV-2 were discovered, the virus began to mutate, and other variants began to develop across the world [6–8]. The B.1.617.2 (Delta) variant first emerged in India in late 2020/early 2021 and rapidly spread to the United Kingdom before spreading to the United States and to 60 other countries across the world [6,9,10]. The SARS-CoV-2 Delta variant is more than twice as infectious as previous variants that developed in 2020 and also caused more than twice as many hospitalizations as the B.1.1.7 (Alpha) variant [10,11].

At the start of the COVID-19 pandemic in 2020, there was a spike in COVID-19 cases in meat packaging plants which caused many of them to temporarily close [12–14]. This could have been largely due to several environmental factors in the meat packaging plants which include air circulation of the virus via HVAC systems, the close proximity of the workers, shared equipment and workspaces, shared travel and living conditions amongst the workers, and the ability of the virus to cohabitate with other biological organisms, like environmental biofilms, which are commonly found in meat packaging plants [15,16].

Biofilms in meat packaging plants are a major threat to food safety, as they are one of the main carriers of foodborne pathogens [17,18]. Biofilms are organized, multicellular assemblages of prokaryotic and eukaryotic cells that are enclosed in a polysaccharide matrix [19]. Biofilms can form on solid, slick surfaces such as tile flooring, PVC pipe, or on stainless steel (SS) [20–23]. Alternatively, biofilms can form on undisturbed water sources such as the inside of drains, puddles, ponds, and lakes [22,24–27]. Bacterial and fungal biofilms have so far been the focus of biofilm research in meat packaging plants [22,28–30]. However, research on the presence of virus particles in the mixed-species biofilm community is still sparse [31–33].

There are several factors to consider when thinking about why biofilms could be an ideal site to harbor SARS-CoV-2 in meat packaging plants. The temperature inside of meat packaging plants is maintained at 4-7°C [12,13]. SARS-CoV-2 virions are stable at colder temperatures and have been shown to persist for several days on materials commonly found in meat packaging plants such as stainless steel, copper, plastic, PVC, and cardboard [34]. Therefore, these facilities have a high risk of harboring and transmitting SARS-CoV-2 [35]. Although bacteria can not directly support virus infection, they can promote viral fitness [33,36,37]. Specifically, some viruses use components of the bacterial envelope to enhance their stability [36,38,39]. Moreover, bacterial communities and biofilms can impact the infection of mammals by viruses [33,36,40]. Furthermore, from a biophysics perspective, virual stability could also be enhanced by the thin liquid film produced by bacterial biofilms [34,41].

There is a critical gap in knowledge in understanding the stability and infectious state of SARS-CoV-2 in multi-species biofilms, particularly those present in meat packaging plants. In this study, SARS-CoV-2 Delta variant was inoculated with- and without three different meat packaging plant environmental biofilms and incubated on SS, PVC, and ceramic tile chips at 7°C. RT-qPCR was used to identify the presence of SARS-CoV-2 Delta variant during incubation, and survival was analyzed by plaque assays to assess the viability of SARS-CoV-2 Delta variant on different surfaces with- and without environmental biofilms, as well as the effect viral presence had on biofilm biomass.

Moreover, the viability of SARS-CoV-2 Delta variant is also linked to the availability of aqueous environments, such as wet surfaces in food processing facilities. Therefore, we performed an analysis to determine how long water takes to evaporate from typical substrate materials found in food processing facilities.

Together, these results indicate that SARS-CoV-2 Delta variant can remain viable and spread throughout meat packaging plants.

## Results

### Mixed-species biofilm cell numbers from all three different meat packaging plants increased in the presence of SARS-CoV-2 Delta variant, on all surfaces tested

To determine if SARS-CoV-2 influences or hinders the growth of an environmental biofilm we grew three different biofilms that consisted of different bacterial populations found in meat packaging plant drains with and without SARS-CoV-2 Delta variant on SS (Fig. 1A), PVC (Fig. 1B) and on ceramic tile chips (Fig. 1C) and incubated for five days at 7°C. The overall mean biofilm cell numbers were represented as colony forming units per mL (CFU/mL).

**Fig. 1.**
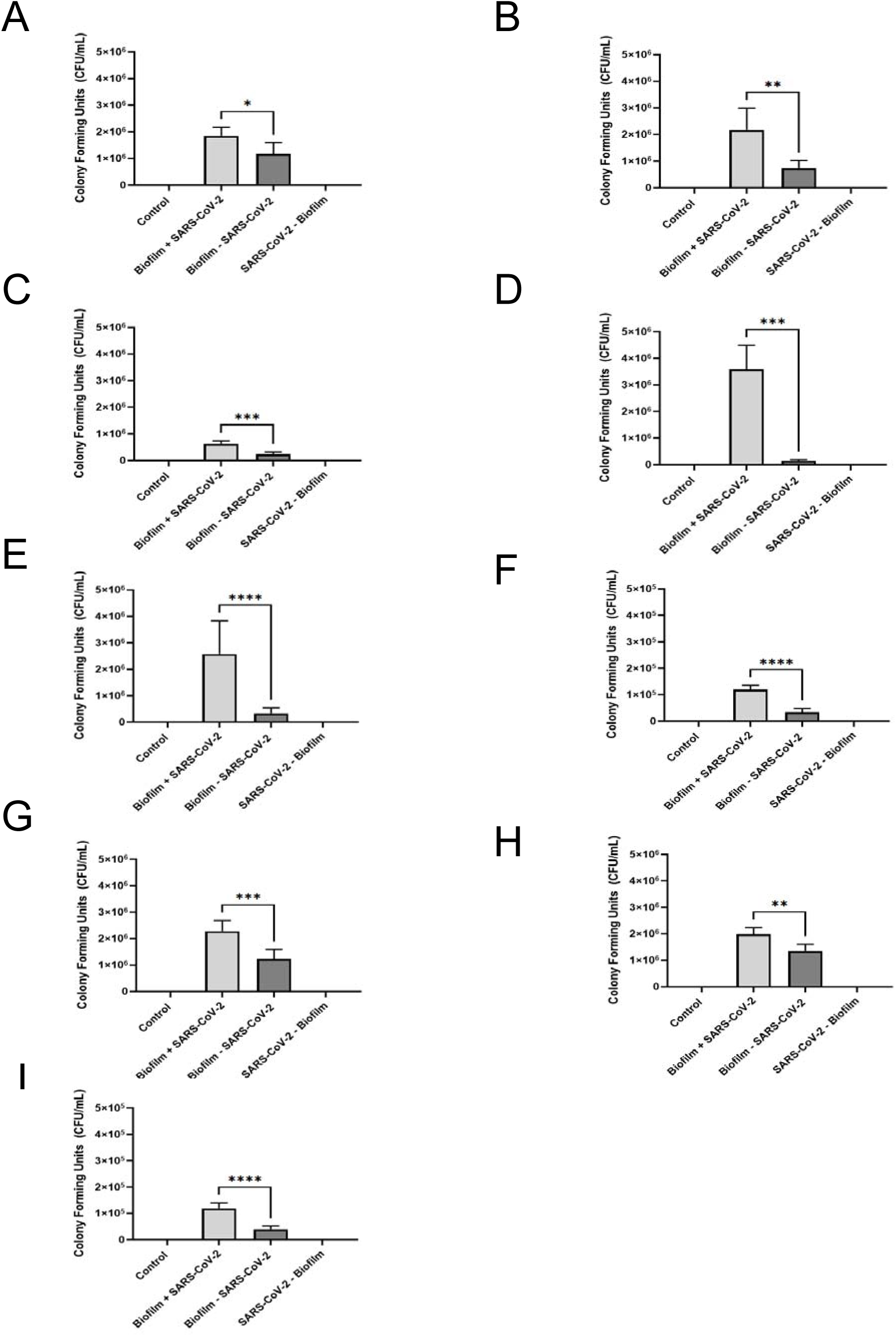
CFU counts from biofilm with SARS-CoV-2 Delta variant and biofilm without SARS-CoV-2 Delta variant samples on stainless steel, PVC, and tile chips. (A-I) CFU counts for biofilm with SARS-CoV-2 Delta variant and biofilm without SARS-CoV-2 Delta variant samples on stainless steel, PVC, and tile chips (A-C) from Plant A, (D-F) from Plant B, and (G-I) from Plant C. Each sample was plated in duplicate. Results in this figure are the mean values and standard deviations (error bars) from three independent experiments. Statistical significance was analyzed by unpaired t-test. *: *p* < 0.05; **: *p* < 0.01; ***: *p* < 0.001; ****: *p* < 0.0001.

Plant A, B, and C bacteria recovered from the biofilms grown in the absence of SARS-CoV-2 Delta variant on SS chips ranged from 1.1 × 10^5^ to 2.0 × 10^6^ CFU/mL, however in the presence of SARS-CoV-2 Delta variant Biofilm A, B, and C numbers were 5.2 × 10^5^ to 3.5 × 10^6^ CFU/mL (Fig. 1A, 1D, 1G). Thus a 1.58-fold increase in the biovolume in the presence of SARS-CoV-2 Delta variant on SS with biofilm organisms from Plant A, a 2.93-fold increase from Plant B, and a 2.65-fold increase from Plant C when compared to the corresponding biofilms grown on SS in the absence of SARS-CoV-2 Delta variant.

The bacterial numbers for biofilm from plants A, B, and C grown in the absence of SARS-CoV-2 Delta variant on PVC chips ranged from 2.0 × 10^4^ to 5.3 × 10^5^ CFU/mL, whereas the number when exposed to SARS-CoV-2 Delta variant on PVC chips ranged from 1.0 × 10^5^ to 4.4 × 10^6^ CFU/mL (Fig. 1B, 1E, 1H); corresponding to a 24.69-fold increase in biovolume for organisms obtained from Plant A on PVC with SARS-CoV-2 Delta variant, a 3.09-fold increase with those from Plant B, and a 3.44-fold increase with Plant C when compared to those obtained in the absence of SARS-CoV-2 Delta variant.

For the biofilms grown on tile chips without SARS-CoV-2 Delta variant the CFU/mL ranged from 2.10 × 10^4^ to 1.6 × 10^6^; whereas in the presence of SARS-CoV-2 Delta variant ranged from 1.0 × 10^5^ to 2.9 × 10^6^ CFU/mL (Fig. 1C, 1F, 1I), representing a 1.86-fold increase in the biofilm obtained from Plant A with SARS-CoV-2 Delta variant, a 1.47-fold increase with those from Plant B, and a 3.04-fold increase with those from Plant C, compared with the biovolumes without SARS-CoV-2 Delta variant. Therefore, our data indicate that SARS-CoV-2 Delta variant positively influences the growth of environmental microorganisms from all three meat packaging plants on all three surfaces: SS, PVC, and ceramic tile.

### RNA levels were lower in the presence of Biofilm B, but had no significant difference for Biofilm A and C

To determine whether meat packaging plant biofilms provide a conducive environment for SARS-CoV-2 Delta, we performed RT-qPCR analyses targeting the nucleocapsid gene (N) of SARS-CoV-2 Delta variant on the harvested samples from biofilms from Plant A, B, and C grown on SS, PVC, and ceramic tile chips, with and without SARS-CoV-2 Delta. We also tested SARS-CoV-2 Delta variant without biofilm organisms on the same surfaces and same incubation conditions. Our RT-qPCR data revealed that there was no statistical significance in the persistence of SARS-CoV-2 Delta variant RNA when a mixed species biofilm from Plant A or C was present (Fig. 2A-2C and Fig. 2G-2I), however, we did detect a significant decrease in N-gene copy number when SARS-CoV-2 Delta variant was mixed with an environmental biofilm organisms from Plant B on all of the materials tested (Fig. 2D-2F).

**Fig. 2.**
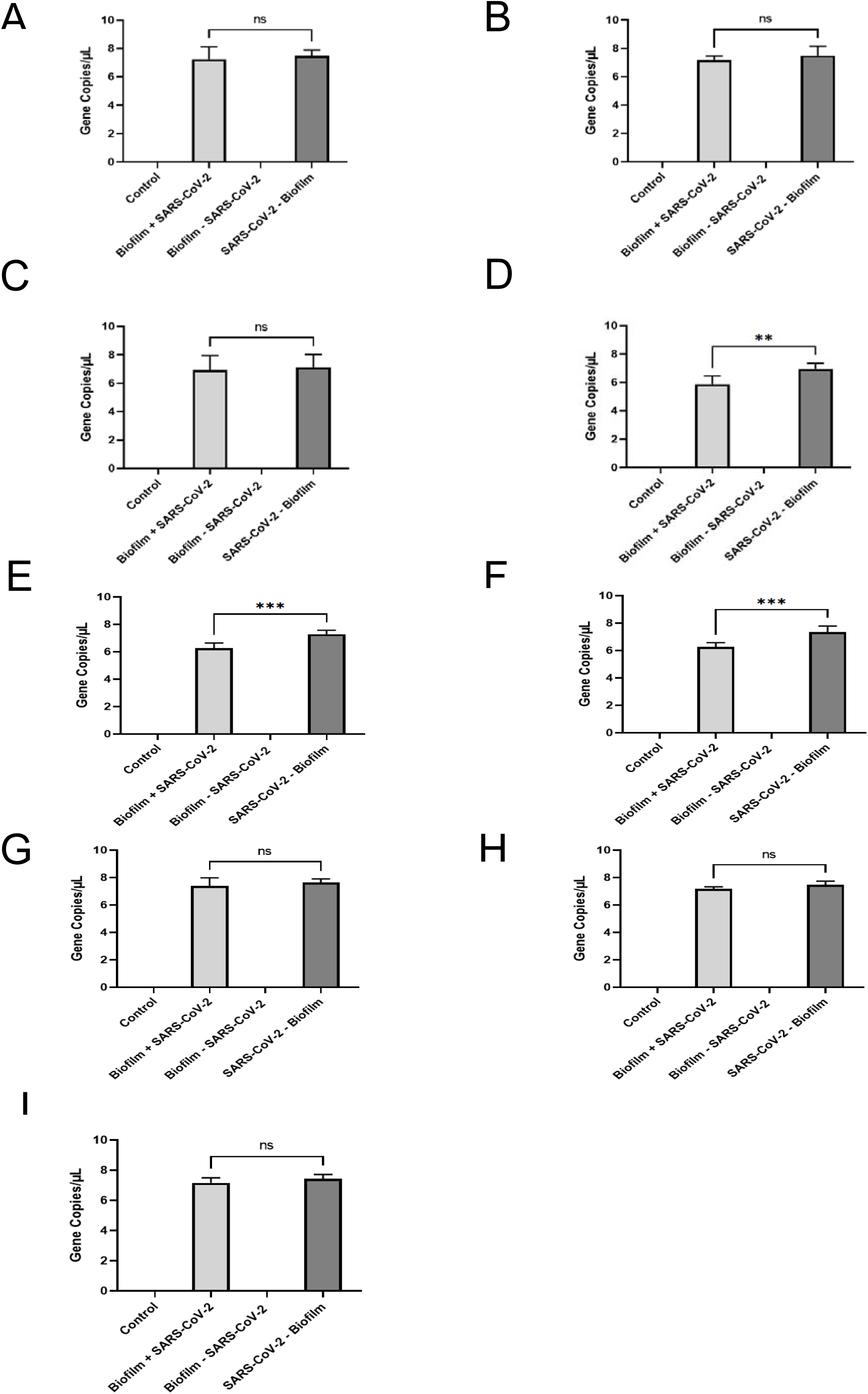
RT-qPCR analysis of SARS-CoV-2 Delta variant mixed with biofilm organisms and pre-incubated for 5 days on stainless steel, PVC, and ceramic tile chips. (A-C) RT-qPCR analysis of SARS-CoV-2 Delta variant mixed with environmental biofilm organisms from Plant A on stainless steel, PVC and on ceramic tile chips, (D-F) RT-qPCR analysis of SARS-CoV-2 Delta variant mixed with environmental biofilm organisms from Plant B on stainless steel, PVC, and on ceramic tile chips, (G-I) RT-qPCR analysis of SARS-CoV-2 Delta variant mixed with environmental biofilm organisms from Plant C on stainless steel, PVC, and on ceramic tile chips. 1.0 × 10^4^ PFU of SARS-CoV-2 Delta variant were added to a stainless steel, PVC, or ceramic tile chip along with a floor drain biofilm sample collected from the cooler of meat packaging plant A, B, or C. The RT-qPCR samples were analyzed in duplicate. Gene copy numbers were calculated from a standard curve of known quantities of SARS-CoV-2 Delta variant RNA in a 25 µL qPCR reaction. Results in this figure are the mean values and standard deviations (error bars) from three independent experiments. Statistical significance was analyzed by unpaired t-test. ns: not significant; **: *p* < 0.01; ***: *p* < 0.001.

When tested on SS the average gene copy numbers for the N-gene for in the presence of SARS-CoV-2 Delta variant was 7.24 gene copies/µL for Plant A, 5.87 gene copies/µL for Plant B, and 7.38 gene copies/µL for Plant C, whereas for SARS-CoV-2 Delta variant by itself on SS chips was 7.47 gene copies/µL for Biofilm A, 6.96 gene copies/µL for Biofilm B, and 7.65 gene copies/µL for Biofilm C (Fig. 2A, 2D, and 2G).

The average gene copy numbers for the N-gene for biofilms grown with SARS-CoV-2 Delta variant on PVC chips was 7.19 gene copies/µL for Plant A, 6.27 gene copies/µL for Plant B, and 7.18 gene copies/µL for Plant C, whereas the gene copy numbers for SARS-CoV-2 Delta variant – Biofilm on PVC was 7.49 gene copies/µL for Biofilm A, 7.29 gene copies/µL for Biofilm B, and 7.47 gene copies/µL for Biofilm C (Fig. 2B, 2E, and 2H).

The average N-gene copy numbers for biofilms grown in the presence of SARS-CoV-2 Delta variant on ceramic tile chips was 6.93 gene copies/µL for Plant A, 6.27 gene copies/µL for Plant B, and 7.16 gene copies/µL for Plant C, whereas the gene copy numbers for SARS-CoV-2 Delta variant – Biofilm on ceramic tile chips was 7.12 gene copies/µL for Biofilm A, 7.34 gene copies/µL for Biofilm B, and 7.44 gene copies/µL for Biofilm C. These results indicate that SARS-CoV-2 Delta variant RNA was significantly degraded when mixed with environmental biofilm organism from Plant B on SS, PVC, and ceramic tile chips compared to when SARS-CoV-2 Delta variant was exposed to SS, PVC, and ceramic tile chips in the absence of biofilm. However, we did not detect a significant reduction in the SARS-CoV-2 Delta variant RNA when inoculated with Biofilm A and C on SS, PVC, and on ceramic tile chips.

### SARS-CoV-2 Delta variant survival was significantly inhibited in the presence of Biofilm B on all surface materials and on Biofilm C on SS and PVC chips

Whilst RT-qPCR analyses is a useful method to identify the presence of an RNA gene target quantitatively, it does not provide any information on the viability of the virus. Therefore, to identify whether SARS-CoV-2 Delta variant was able to survive and remain infectious when incubated with environmental biofilms, we performed plaque assays to quantitatively analyze the number of infectious virus particles recovered after incubation on the three different surface materials that we tested in this study with and without an environmental biofilm. In short, 1 × 10^4^ virus particles were inoculated onto surface materials with and without environmental biofilm organisms, and viral infectivity was measured using a solid double overlay plaque assay. For all the materials tested, a significantly lower average plaque forming units (PFU)/mL was detected in the presence of biofilm organisms from Plant B. Lower average PFUs were also observed when the virus was incubated with the biofilm organism from Plant C on SS and PVC chips (Fig. 3D - 3H).

**Fig. 3.**
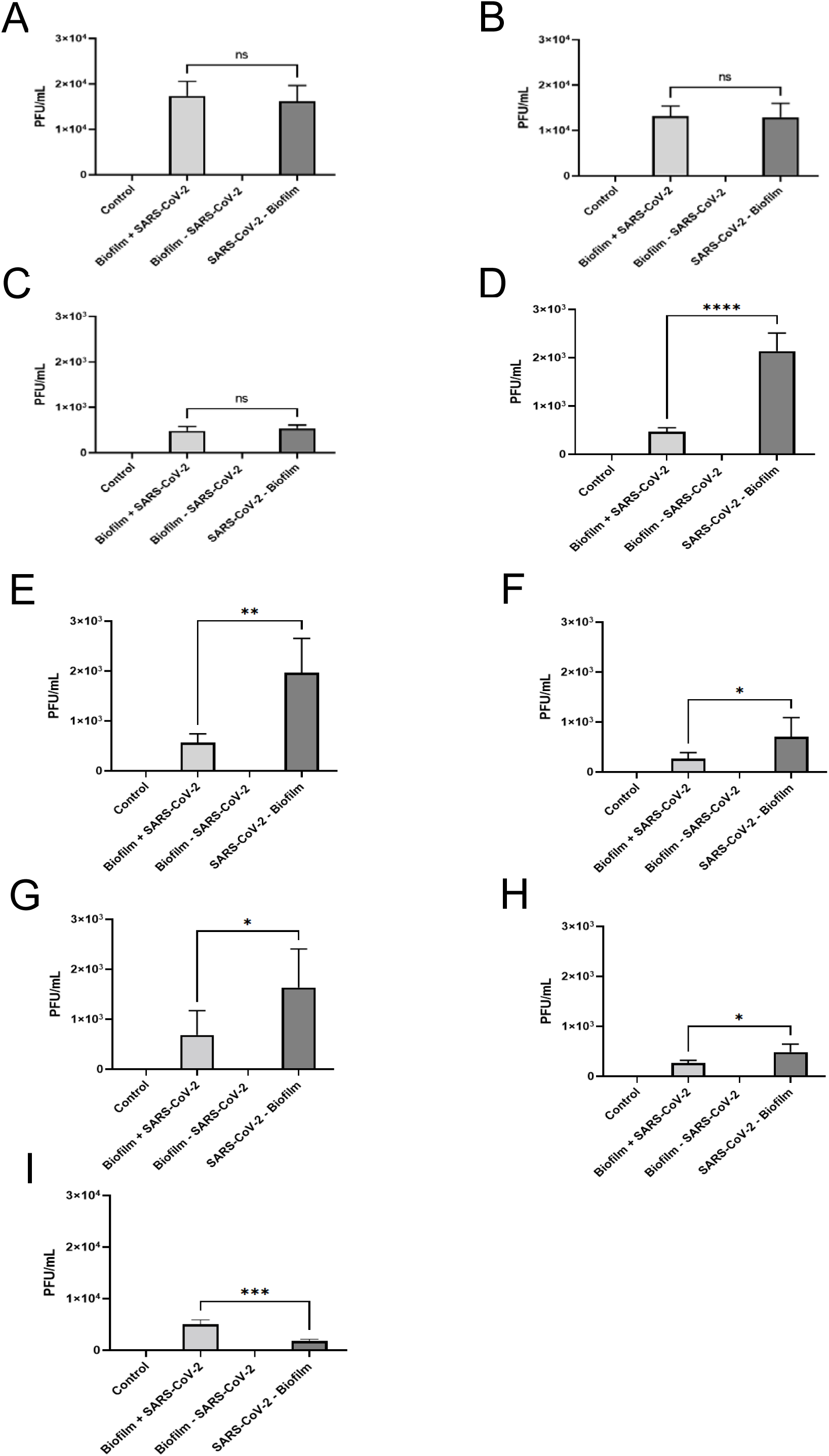
Plaque assay results from biofilm with SARS-CoV-2 Delta variant and SARS-CoV-2 Delta variant without biofilm samples on stainless steel, PVC, and ceramic tile chips. (A-I) Results from plaque assays on samples collected from (A-C) stainless steel, (D-F) PVC, and (G-I) ceramic tile chips. Each sample was filtered through a 0.45 µm filter and plated on Vero CCL-81 cells in duplicate. Results in this figure are the mean values and standard deviations (error bars) from three independent experiments. Statistical significance was analyzed by unpaired t-test. **: *p* < 0.01; ****: *p* < 0.0001.

The average PFU/mL for SARS-CoV-2 Delta variant incubated with biofilm organism on SS chips was 1.73 × 10^4^ PFU/mL for Plant A, 4.67 × 10^3^ PFU/mL for Plant B, and 683 PFU/mL for Plant C, whereas the average PFU/mL for Biofilm – SARS-CoV-2 Delta variant on SS chips was 16,167 PFU/mL for Biofilm A, 21,333 PFU/mL for Biofilm B, and 1,633 PFU/mL for Biofilm C (Fig. 3A, 3D, and 3G). For SS chips there was no significant difference between the PFU/mL for SARS-CoV-2 incubated to- or without biofilm organisms from Plant A, however, there was a 4.57-fold reduction in infectious SARS-CoV-2 Delta variant when exposed to biofilm organism from Plant B, and a 2.39-fold reduction in infectious SARS-CoV-2 Delta variant after exposure to biofilm organisms from Plant C.

Similarly, SARS-CoV-2 Delta variant on exposed to biofilm on PVC chips gave 1.32 × 10^5^ PFU/mL for Plant A, 5.67 × 10^3^ PFU/mL for Plant B, and 267 PFU/mL for Biofilm C, whereas the average PFU/mL for SARS-CoV-2 Delta variant - Biofilm was 128,333 PFU/mL for Biofilm A, 19,667 PFU/mL for Biofilm B, and 483 PFU/mL for Biofilm C (Fig 3B, 3E, and 3H). For PVC chips there was no significant difference between the PFU/mL for SARS-CoV-2 Delta variant exposed to- and without biofilm forming organisms from Plant A, however, there was a 3.47-fold reduction in infectious SARS-CoV-2 Delta variant when exposed to the biofilm organisms from Plant B, and a 1.81-fold reduction in PFU when incubated with organism from Plant C.

When incubated on tile chips the average PFU/mL for SARS-CoV-2 Delta variant exposed to biofilm forming organisms was 4.83 × 10^3^ PFU/mL for Plant A, 267 PFU/mL for Plant B, and 5 × 10^3^ PFU/mL for Plant C, whereas the average PFU/mL for SARS-CoV-2 Delta variant – Biofilm was 5,333 PFU/mL for Biofilm A, 483 PFU/mL for Biofilm B, and 1,817 PFU/mL for Biofilm C (Fig 3C, 3F, and 3I). For the ceramic tile chips there was again no significant difference between the PFU/mL for SARS-CoV-2 Delta variant incubated with- and without biofilm forming organism from Plant A, however, there was a 2.62-fold reduction in PFU/mL after exposure to the biofilm forming organisms from Plant B, and a 2.75-fold increase in infectious SARS-CoV-2 Delta after exposure to the biofilm organisms from Plant C, compared with virus incubated alone. These results indicate that SARS-CoV-2 Delta variant had no significant reduction in infectivity when mixed with organisms from Plant A when incubated on all the test materials. However, there was a significant reduction in infectivity of SARS-CoV-2 Delta variant when exposed to the biofilm forming organisms from Plant B on all of the materials tested. Plant C organisms showed a significant effect on reducing SARS-CoV-2 Delta variant infectivity when incubated on SS and PVC chips, but was able to offer SARS-CoV-2 protection when incubated on tile chips, so that viability was higher than that obtained when the virus was incubated by itself.

### Evaporation dynamics for different substrate materials in meat processing facilities

To examine the availability of aqueous environments for SARS-CoV-2 Delta variant to survive in meat processing facilities, we performed an analysis to determine how long liquid takes to evaporate from typical substrates found in such facilities. We measured the evaporation rates from stainless steel (red, circles), PVC (green, diamonds) and ceramic tile (blue, triangles) samples. Fig 5(A) shows the weight-fraction of liquid remaining on each of these substrates as a function of time (hours) post inoculation. These data points suggest that water evaporates faster from stainless steel compared to PVC and ceramic tiles. To quantify this, we performed a least-square fitting analysis to an exponential decay function to determine the half-life time of the liquid on each of these substrates. Fig 5(B) shows these half-life times, giving 88 ± 9 hours for stainless steel, 110 ± 16 hours for PVC, and 127 ± 10 hours for ceramic tile. Thus, the PVC and ceramic tiles provide a more stable aqueous environment for SARS-CoV-2 Delta variant to remain viable longer compared to stainless steel.

## Discussion

At the start of the SARS-CoV-2 pandemic in 2020, many meat packaging plants had to be closed due to the high number of SARS-CoV-2 cases amongst the workers [13,42–44]. These closures created a bottleneck in the supply chain between the livestock producers, feedlot operators, and the processors. To determine why SARS-CoV-2 had a high occurrence rate in meat packaging plants, we investigated if SARS-CoV-2 Delta variant could survive within meat packaging plant biofilms as a potential mechanism for SARS-CoV-2 endurance and persistence. We demonstrated in this study that SARS-CoV-2 Delta variant was able to remain viable for up to five days post-inoculation on SS, PVC, and on tile chips with- and without environmental biofilms from three different meat packaging plants. Therefore, meat packaging plants are at a high risk of harboring SARS-CoV-2 and spreading the virus amongst the workers in these facilities.

In addition to meat packaging plants being a conducive environment for SARS-CoV-2 to survive and disseminate, meat packaging plants are also an opportune environment for environmental biofilms. Environmental biofilms in meat packaging plants can be a source of foodborne pathogen outbreaks that are a serious threat to food safety and human health [20,22]. Biofilms can develop on a wide range of diverse surfaces throughout the meat packaging plant such as floors, drains, and areas that are hard to reach and do not come into contact with surface sanitizer very often [24,45]. The protective matrix of the biofilm can also offer shelter to organisms within the biofilm from the effects of disinfecting agents.

Biofilms have been suggested to act as a reservoir for the survival and spread of other viruses, such as noroviruses [32,33,36,46,47]. In this study, we utilized floor drain samples that were collected from different meat packaging plants to identify the viability of SARS-CoV-2 Delta variant with- and without an environmental biofilm on several common materials found in meat packaging plants: SS, PVC, and tile chips. We observed that SARS-CoV-2 Delta variant can remain not just detectable but also viable on all of the materials tested (Fig. 2 & Fig. 3). We also observed that the viability of the virus on each material tested was not significantly different with and without biofilm forming organisms from Plant A, however, the viability of the virus was reduced in the presence of organisms from Plant B on each material tested and for Plant C on stainless steel and on PVC chips (Fig. 2 & Fig. 3).

The viability of SARS-CoV-2 in the environmental biofilm was detected via a solid double overlay plaque assay to identify the number of infectious virus particles and via RT-qPCR to quantifiably detect viral RNA on each material tested compared to SARS-CoV-2 Delta variant inoculated on the materials by itself (Fig. 2 and Fig. 3). The RT-qPCR data suggests that most of the SARS-CoV-2 Delta variant mixed with biofilm was from non-viable or inactive virus since the plaque assay data differed considerably (Fig. 2 and Fig. 3). The PFU/mL for SARS-CoV-2 Delta variant – Biofilm B or C on stainless steel or PVC chips indicated a 46.88 – 207.04-fold reduction of infectious SARS-CoV-2 Delta variant virus particles compared to the initial titer of the virus (1.0 × 10^5^ PFU/mL) after incubating on the different materials for five days at 7°C. However, when SARS-CoV-2 Delta variant was mixed with biofilm forming organisms from Plants B and C and inoculated on SS or PVC chips, we observed a 146.41 – 374.53-fold reduction in infectivity compared to the original titer of the virus (1.0 × 10^5^ PFU/mL). We did not detect a significant reduction in infectivity for SARS-CoV-2 Delta variant exposed to organisms from Plant A on SS, PVC, or on ceramic tile chips.

Our plaque assay studies indicated that SARS-CoV-2 Delta variant was able to remain infectious when mixed with biofilm forming organism from Plants A, B, and C on SS, PVC, and on ceramic tile chips for five days at 7°C (Fig. 3). SARS-CoV-2 Delta variant was more infectious following incubation on SS and PVC chips after exposure to biofilm forming organisms from Plants A & B, compared the equivalent PFUs after incubation on tile chips. After exposure to the biofilm forming organisms from Plant C, SARS-CoV-2 Delta variant was more infectious on tile chips than SS, and then more viable on SS when compared to PVC chips. Interestingly, SARS-CoV-2 Delta variant showed higher viability after exposure to the biofilm organisms from Plant C on tile chips, than when it was incubated on tile chip alone. Even a modest amount of virus survival in the meat packaging plant could be a health risk for the transmission and spread of SARS-CoV-2 within the meat packaging plant.

These results suggest that the viability of SARS-CoV-2 Delta variant is highly dependent on the microorganisms that are present within each biofilm. Previous work on the population structures for the biofilms obtained from the different plants has shown that Plant A is composed of xxx….

One of the most surprising results from our study was the increase in biofilm biovolume for all three biofilms in the presence of SARS-CoV-2 Delta variant, ranging from a 1.47 – 24.69-fold increase compared to when the biofilms were inoculated without virus on all of the materials tested (Fig. 1). These results correlate with what others have previously shown, in that virus particles can enhance the biovolume of the biofilm [32,33,46]. In nature, bacteria interact with eukaryotes and other prokaryotes, fighting for survival through synergistic, mutualistic, and antagonistic interactions [18,32,33,36,46]. The increase in biovolume could be linked to the virus triggering a defense mechanism in the bacteria which results in the bacteria increasing their biovolume so it can expand in the presence of the virus (Fig. 1). This result is critical in elucidating the interactions between the bacteria and virus and how they can potentially work together and react to one another.

In the evaporation dynamics assays, we examined how the viability of SARS-CoV-2 Delta variant is linked to the availability of aqueous environments, such as wet surfaces in food processing facilities. We quantified how long liquid can be retained on commonly found substrate materials in food processing facilities, including stainless steel, PVC, and ceramic tiles. Our results indicate that water evaporates faster from stainless steel compared to PVC and tile chips. Hence, the latter two provide more favorable conditions for the virus.

Our results indicate that SARS-CoV-2 Delta variant can remain viable for up to five days within each biofilm from three different meat packaging plants. Our conclusions suggest that SARS-CoV-2 could easily spread among the workers in the meat packaging plant, remaining viable on SS, PVC, and on ceramic tile chips. We observed that SARS-CoV-2 Delta variant was, for the most part, more viable in the absence of biofilm, but was able to remain infectious in the presence of biofilm. Our data suggests that wash-water carrying SARS-CoV-2 could seep into the drains of the meat packaging plant and the virus remain viable in the drainage system, interacting with biofilm forming organisms. Biofilms could potentially facilitate the survival of SARS-CoV-2 throughout the facility through several active processes, such as the virus binding to the biofilm polysaccharide matrix, preventing desiccation and exposure to sanitizing agents. The biofilm could also help to spread the virus through bacterial motility; the bacteria can also undergo swarming, which would allow the virus to potentially move outwards as the biofilm develops new extracellular matrices, spreading across the meat packaging plant through the drain systems [22,33,48,49]. It is well documented that SARS-CoV-2 can spread through aerosols [2,3,5,50]. In addition, another method for how SARS-CoV-2 can spread throughout the meat packaging plant would be from when the floors are washed with high-pressure water. When the high-pressured water hits the drain system, it is possible that the water can disrupt the biofilm containing the virus creating aerosols that then spread throughout the facility. Once airborne, the HVAC system in the meat packaging plant and the colder temperature can facilitate the survival and distribution of SARS-CoV-2 throughout the facility, which is in agreement with current fluid mechanic models [51].

Our findings in this study have led us to conclude that the viability of the SARS-CoV-2 Delta variant with and without the biofilms, the survival on surfaces at 7°C, current models of spread via HVAC systems, and fluid mechanics models all provide evidence for the long-term survival and spread of SARS-CoV-2 in meat packaging plants. These findings, along with the close working quarters, shared equipment, and shared travel to and from the meat packaging plant all provide an environment where SARS-CoV-2 can be rapidly transmitted. Continued studies on the survival and dispersal of SARS-CoV-2 in meat packaging plants will hopefully provide new details that can inform and help reduce the spread of SARS-CoV-2 and other pathogens within these environments.

## Conclusions

Our data provides evidence that SARS-CoV-2 Delta variant can persist and remain viable with- and without environmental biofilms found in meat packaging plants under the typical environmental conditions found in meat packaging plants. We identified a difference in viral viability that was dependent on the microbial structure and make-up of the biofilms tested. These results suggest that biofilms could act as a reservoir for SARS-CoV-2 Delta variant to persist and spread throughout meat packaging plants. The results from this study provide evidence for why high numbers of cases of COVID-19 have occurred in in meat packaging plants. Our CFU results provide evidence that SARS-CoV-2 Delta variant stimulates the bacteria found in the environmental biofilms, resulting in an increase to their biovolumes. Future work will need to be conducted to understand the biological interactions between the virus and biofilm, such as the protein-protein interactions between the virus and bacteria, positional virus survival within the biofilm, bacterial quorum sensing of the virus, and transcriptomics within each biofilm population to understand which genes are being upregulated and downregulated, and if different species respond in different manners, in the presence of SARS-CoV-2 Delta variant. Altogether, this work will help in understanding viral and bacterial interactions, allowing for the design of intervention strategies to help prevent future bacterial and viral outbreaks from occurring in meat packaging plants.

## Materials and Methods

### Drain sample collection and characterization

The meat packaging plant floor drain biofilm samples were collected from three different meat processing plants, following the previously described protocol [22] and were generously provided for this study by Drs. Mick Bosilivac and Rong Wang USDA-ARS-USMARC, Clay Center, Nebraska.

### Cell lines and SARS-CoV-2 propagation

Vero CCL-81 cells (ATCC® CCL-81) were used for the propagation of SARS-CoV-2 viral particles and for the solid double overlay plaque assays. Vero CCL-81 cells used in this study were cultured at 37°C in 5% CO_2_ in Dulbecco’s modified Eagle medium (DMEM; Cellgro) supplemented with 10% fetal bovine serum (FBS), penicillin (50 IU/mL), and streptomycin (50 µg/mL). SARS-CoV-2 Delta variant was used for all of the experiments in this study and was acquired from the ATCC (ATCC NR-55672, hCoV-19/USA/MD-HP05647/2021 Delta, batch number: 70046635). The virus stocks used for this study were produced as previously described (31).

### Assay of SARS-CoV-2 infectivity

Viral infectivity was determined by titrating viral stock onto cultured Vero CCL-81 cells and a solid double overlay plaque assay was performed as previously described [52]. The SARS-CoV-2 viral titer used for all experiments was 1.0 × 10^5^ PFU/mL.

For the recovered samples 300 µL of each homogenate was filtered through a 0.45 µm syringe filter to remove bacterial contaminants before being serially diluted in DMEM with 2% FBS and 1% Streptomycin/Penicillin mix. Each sample was plated onto cultured Vero CCL-81 cells in duplicate. Results from this experiment are the mean values and standard deviations (error bars) from three independent experiments. A previously published protocol was followed for the solid double overlay plaque assay [52].

### Biofilm formation with drain sample and SARS-CoV-2

SARS-CoV-2 stocks were cultured to a viral titer of 1.0 × 10^5^ PFU/mL prior to the start of the experiment and stored at −80°C [53]. To simulate the meat packaging plant environment, floor drain samples were 50-fold inoculated into Lennox Broth without salt medium (LB-NS, Acumedia Manufacturers, Baltimore, MD) and incubated at 7°C for 5 days with orbital shaking at 200 rpm [53]. On the fifth day, a 1.0 mL aliquot was removed from each sample, diluted in sterile LB-NS medium, and plated onto Trypticase soy agar (TSA) plates for colony enumeration after overnight incubation at 37 °C. To investigate whether biofilm formation from meat packaging plant floor drain samples can support the harborage of SARS-CoV-2, biofilms with or without SARS-CoV-2 Delta variant on SS, PVC, and ceramic tile (Fig. 3). Controls included SARS-CoV-2 Delta variant alone (no biofilm) and a media only negative control. The experiments were set-up in duplicate in 6-well plates, and repeated three times, to give a total of six data points for each assay condition. Each well contained one sterile (18×18×2mm) SS, PVC, or ceramic tile chip. The following test combinations were added to the top facing surface of each chip: (A) Biofilm with SARS-CoV-2 Delta variant: 100 µL of the 5-day floor drain pre-culture (described above) mixed with 100 µL of SARS-CoV-2 Delta variant in DMEM and 100 µL of LB-NS media; (B) Biofilm without SARS-CoV-2 Delta variant: 100 µL of the five-day biofilm pre-culture, 100 µL of DMEM, and 100 µL of LB-NS media; (C) SARS-CoV-2 Delta variant alone (no biofilm): 100 µL of SARS-CoV-2 Delta variant (1 × 10^4^) in DMEM and 100 µL of LB-NS media; (D) Media only control: 100 µL of DMEM and 200 µL of LB-NS. Each experimental variable was incubated at 7°C for five days.

At the end of the incubation period, biofilm biomass/virus was harvested from each chip by lifting the chip with sterile forceps, scraping the material on both sides with a sterile cell scraper into a sterile tube and rinsing the chip with 1 mL of LB-NS, which was also collected (Fig 4). The collected sample was homogenized by pipetting. The drain biofilm biomass was determined by taking 100 µL of the homogenate, performing 10-fold dilutions into LB-NS and plating on TSA plates for colony enumeration following an overnight incubation at 37°C. The remaining homogenate was used for RT-qPCR and plaque assay analysis. Results from this experiment are the mean values and standard deviations (error bars) from three independent experiments, run in duplicate.

**Fig. 4.**
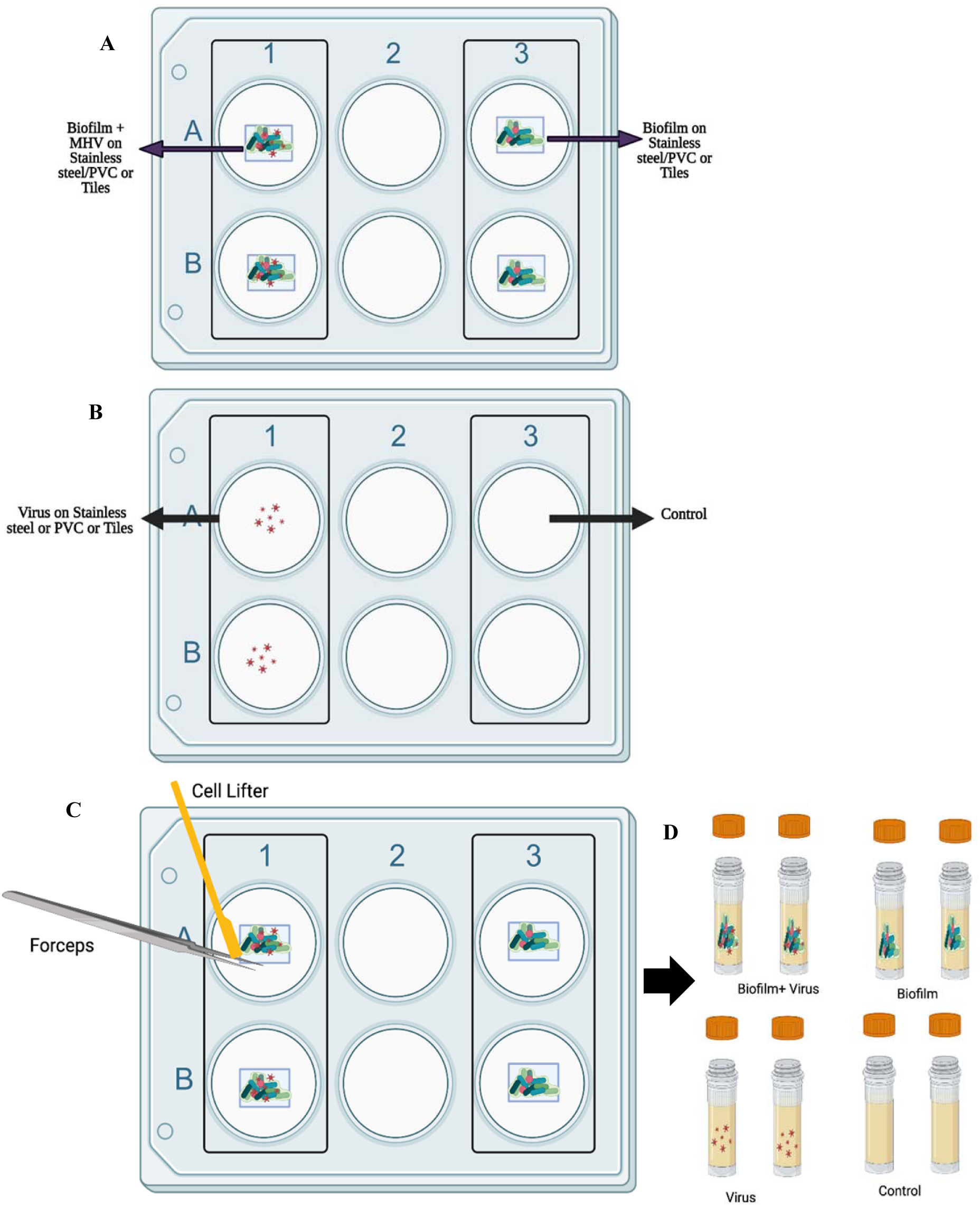
Schematic representation of floor drain biofilm and virus experiment. (A and B): Experimental set up with Biofilm with SARS-CoV-2 Delta variant, Biofilm without SARS-CoV-2 Delta variant, SARS-CoV-2 Delta variant - Biofilm, and Negative Control in duplicate. The experimental set is incubated at 7°C for 5 days. (C). After 5 days, the biofilm was harvested from SS, PVC, or ceramic tile chips using a cell lifter and forceps and rinsed with 1000 µL of LB-NS. (D) Harvested cells were stored in a screw-cap tube at −80°C until needed.

**Fig. 5.**
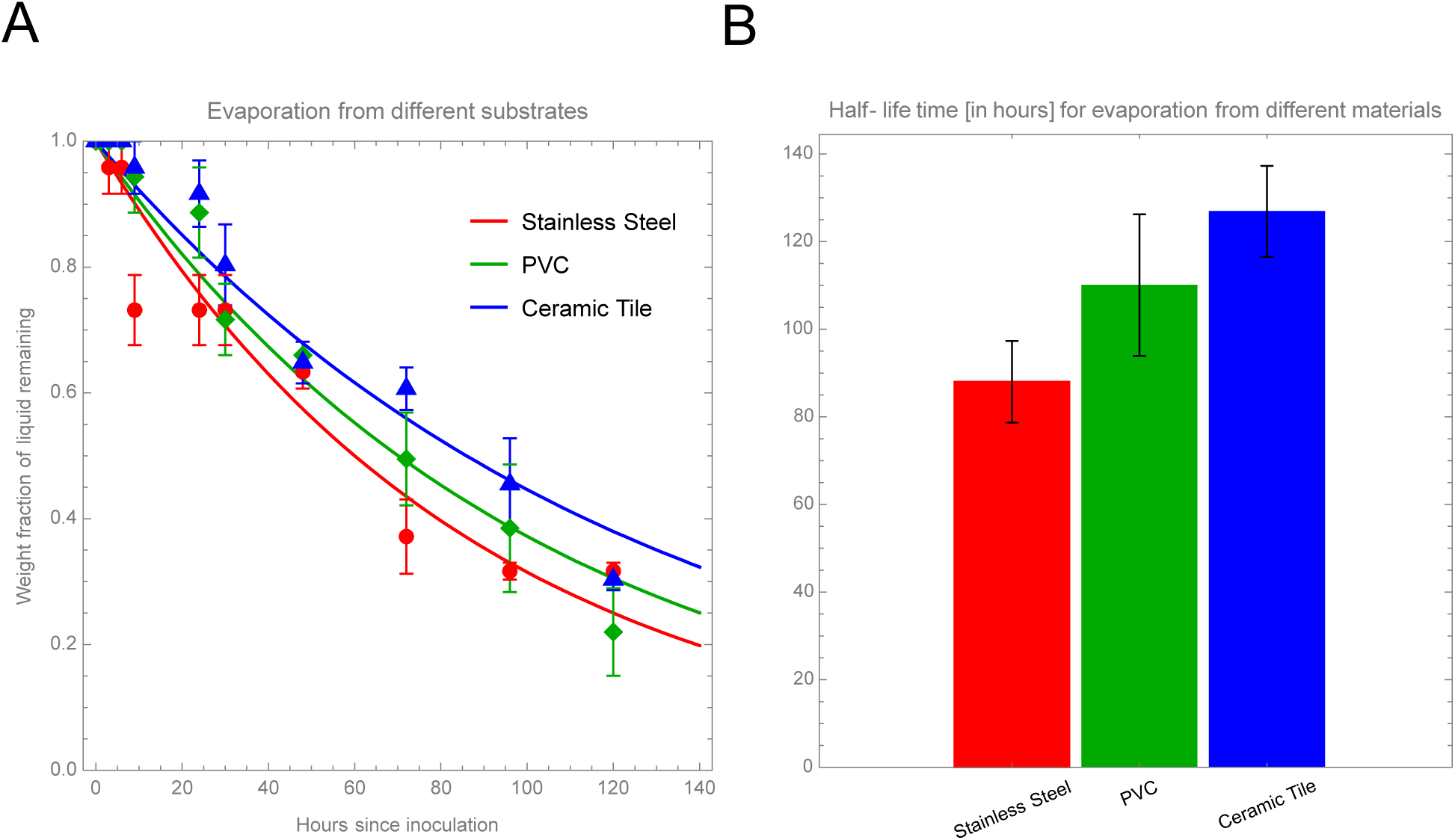
Results from evaporation dynamics assays of water droplets inoculated on different substrates: stainless steel (red, circles), PVC (green, diamonds) and ceramic tile (blue, triangles) samples. (A) Weight fraction of liquid remaining on the substrates as a function of time (hours) after inoculation. The data points represent mean values over N=6 replicates, and the error bars show the standard error (SE) over these replicates. The curves show exponential decay fits to these data points. (B) Half-life time of evaporation from the different materials, obtained from these exponential decay fits. This gives 88 ± 9 hours, 110 ± 16 hours, 127 ± 10 hours, respectively, where the error bars are quantified by the standard error (SE) of the data sets.

### SARS-CoV-2 RT-qPCR Analysis

Viral RNA from each sample was extracted and purified to perform RT-qPCR to determine the relative copy numbers of SARS-CoV-2 Delta variant in each sample. Viral RNA was extracted and purified using Zymo’s Quick-DNA/RNA Viral Magbead Extraction kit along with a Thermo Scientific Kingfisher Flex machine. Purified RNA samples were quantified by using a SpectroStar Nano spectrophotometer. Purified RNA samples were stored at −20°C. Taqman-based RT-qPCR analyses were completed using NEB’s Luna^®^ Universal Probe One-Step RT-qPCR kit. Purified RNA extracted from SARS-CoV-2 Delta variant was used for the positive control and to create a standard curve. The RT-qPCR reactions were completed in 25 µL volumes using the Luna Universal Probe One-Step Reaction Mix. The RT-qPCR mixture contained 10 µL of Luna Universal Probe One-Step Reaction Mix, 1 µL of Luna WarmStart RT Enzyme Mix, 400 nM of nCOV_N1 Forward Primer (IDT Catalog #10006821), 400 nM of nCOV_N1 Reverse Primer (IDT Catalog #10006822), 200 nM of nCOV_N1 probe (IDT Catalog #10006823), 250 ng RNA, and nuclease free water. The RT-qPCR analysis was performed using a Bio-Rad CFX96 Deep Well Real Time thermal cycler. Reverse transcription occurred at 55°C for 10 minutes, after which there was denaturation and *Taq* polymerase activation at 95°C for 1 minute, and then 40 cycles at 95°C for 15 seconds followed by 60°C for 30 seconds for data collection. RT-qPCR reactions were performed in duplicate for each sample and the sample threshold cycle (CT) was used for data analysis. Gene copy numbers were calculated by comparing the CT value for 250 ng SARS-CoV-2 Delta variant on the standard curve, with the CT value for each sample. The following equation was used to calculate the gene copy numbers for the N-gene of SARS-CoV-2 Delta variant: Gene Copy Number = (Copy Number of 250 ng of positive control) - ((CT Pos Cont. - CT exp cont)/CT exp cont)*(Copy number of 250 ng of positive control)[54]. Data from each sample was compared using positive and negative controls performed in duplicate. Results from this experiment are the mean value and standard deviation (error bars) from three independent experiments (Fig. 2).

### Evaporation dynamics assays

We performed an analysis to determine how long liquid takes to evaporate from typical substrates found in meat processing facilities, to examine the availability of aqueous environments for SARS-CoV-2 to survive in these facilities. We measured the evaporation rates from fixed-size samples of stainless steel, PVC, and ceramic tile. These substrates were inoculated with a calibrated amount of culture media, after which the weight of the samples plus the remaining media was measured with a high-precision balance at time points of 0, 1, 3, 6, 9, 24, 30, 48, 72, 96, and 120 hpi. For each substrate type, we performed N=6 replicates. From these measurements, we determine the fraction (percentage/100) of weight of the media remaining on the substrates, compared to the initial weight of the media at 0 hpi. These results are shown in Figure 5. Panel (A) shows this weight fraction, *f*(*t*), as a function of time post inoculation, *t*. The data points and error bars represent the mean and standard error of these replicates, respectively. For each substrate, we then performed a least-squares fit analysis of the data to an exponential decay function, *f*(*t*) = exp(−*t/τ*), where τ is the half-life time of the media. Panel (B) shows these half-life times for the three substrates, where the error bars again represents the standard error of the ensemble of replicates.

**Table 1:**
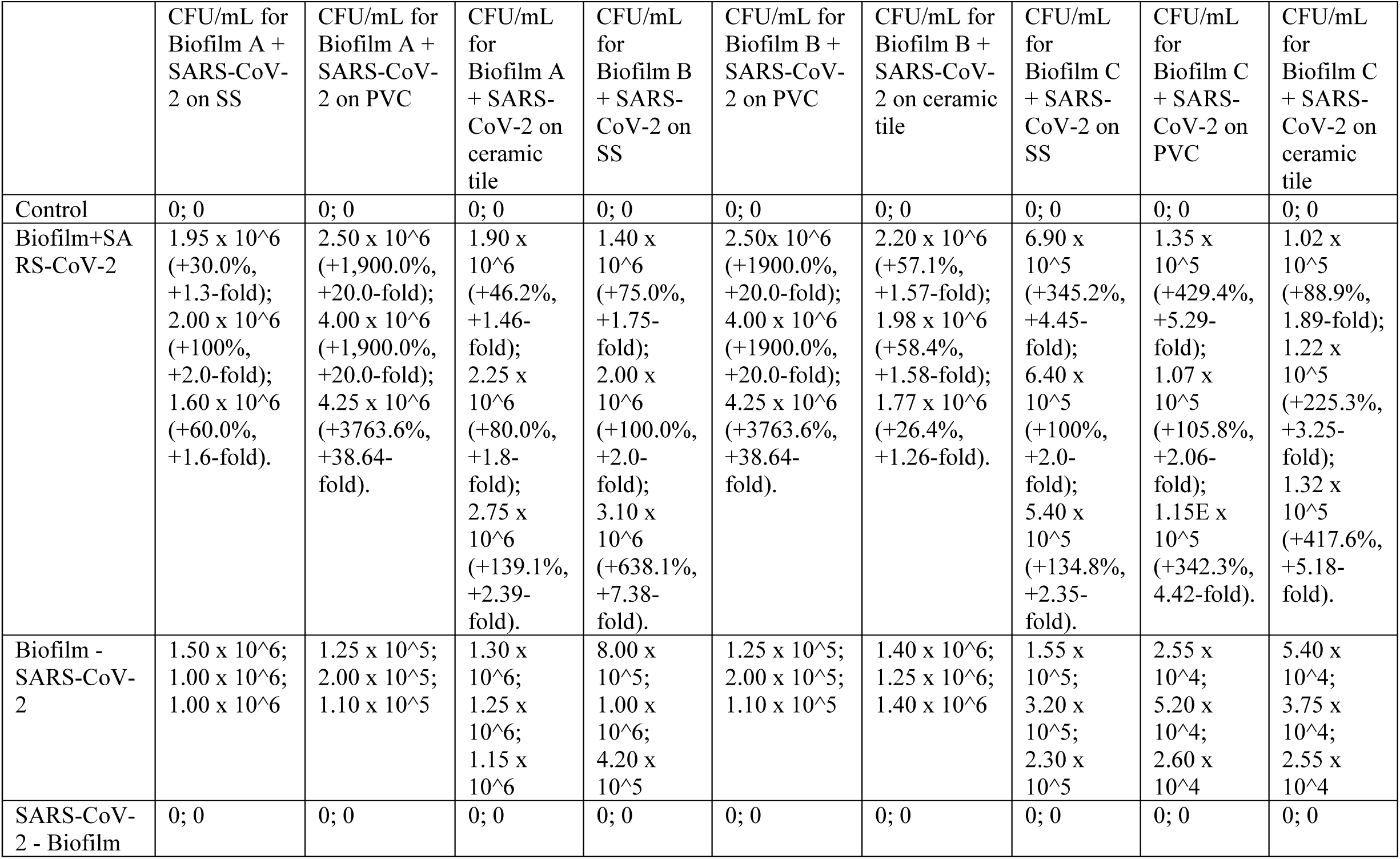
Data from the Biofilm +/− SARS-CoV-2 Delta variant CFU/mL count from different experimental conditions. Table 1 indicates CFU/mL numbers and the percentage and fold change compared to the initial biofilm inoculum.

**Table 2:** Data from the Biofilm +/− SARS-CoV-2 Delta variant RT-qPCR analyses on the recovered SARS-CoV-2 Delta variant RNA from the different experimental conditions. Table 2 indicates CT numbers and the percentage and fold change from the initial inoculum (1.0 × 10^4^).

|  | N-gene CT #<br>for Biofilm A<br>+ SARS-CoV-2 on SS | N-gene<br>CT # for<br>Biofilm A<br>+ SARS-CoV-2 on<br>PVC | N-gene<br>CT # for<br>Biofilm A<br>+ SARS-CoV-2 on<br>ceramic<br>tile | N-gene<br>CT # for<br>Biofilm B<br>+ SARS-CoV-2 on<br>SS | N-gene<br>CT # for<br>Biofilm B<br>+ SARS-CoV-2 on<br>PVC | N-gene<br>CT # for<br>Biofilm B<br>+ SARS-CoV-2 on<br>ceramic<br>tile | N-gene<br>CT # for<br>Biofilm C<br>+ SARS-CoV-2 on<br>SS | N-gene<br>CT # for<br>Biofilm C<br>+ SARS-CoV-2 on<br>PVC | N-gene CT<br># for<br>Biofilm C<br>+ SARS-CoV-2 on<br>ceramic tile |
| --- | --- | --- | --- | --- | --- | --- | --- | --- | --- |
| Control | 0; 0 | 0; 0 | 0; 0 | 0; 0 | 0; 0 | 0; 0 | 0; 0 | 0; 0 | 0; 0 |
| Biofilm+SARS-<br>CoV-2 | 22.9<br>(+13.4%,<br>+1.13-fold);<br>22.0 (+3.3%,<br>+1.03-fold);<br>17.2 (-6.5%, -<br>1.07-fold). | 21.4<br>(+4.4%,<br>+1.04-<br>fold);<br>20.3 (-<br>1.9%, -<br>1.02-fold);<br>20.7<br>(+12.5%,<br>+1.13-<br>fold). | 22.9<br>(+6.5%,<br>+1.07-<br>fold);<br>22.3<br>(+1.4%,<br>+1.01-<br>fold);<br>19.6<br>(+0.0%,<br>+0.0-<br>fold). | 24.9<br>(+16.4%,<br>1.16-fold);<br>23.7<br>(+16.2%,<br>+1.16-<br>fold);<br>21.4<br>(+12.6%,<br>+1.13-<br>fold). | 23.2<br>(+22.8%,<br>+1.23-<br>fold);<br>21.9<br>(+14.1%,<br>+1.14-<br>fold);<br>21.6<br>(+9.7%,<br>+1.10-<br>fold). | 22.5<br>(+15.4%,<br>+1.15-<br>fold);<br>22.4<br>(+12.0%,<br>+1.12-<br>fold);<br>21.8<br>(+21.8%,<br>+1.22-<br>fold). | 19.7<br>(+3.7%,<br>+1.04-<br>fold);<br>20.0<br>(+7.0%,<br>+1.07-<br>fold);<br>18.3<br>(+1.7%,<br>+1.02-<br>fold). | 19.7<br>(+5.9%,<br>+1.06-<br>fold);<br>20.3<br>(+7.4%,<br>1.07-fold);<br>19.8<br>(+0.0%,<br>+0.0-<br>fold). | 19.4<br>(+2.6%,<br>+1.03-<br>fold);<br>21.1<br>(+12.8%,<br>+1.13-<br>fold);<br>19.5 (-<br>2.0%, -<br>1.02-fold). |
| Biofilm -<br>SARS-CoV-2 | 0; 0 | 0; 0 | 0; 0 | 0; 0 | 0; 0 | 0; 0 | 0; 0 | 0; 0 | 0; 0 |
| SARS-CoV-2 -<br>Biofilm | 20.2;<br>21.3;<br>18.4 | 20.5;<br>20.7;<br>18.4 | 21.5;<br>22.0;<br>19.6 | 21.4;<br>20.4;<br>19.0 | 18.9;<br>19.2;<br>19.7 | 19.5;<br>20.0;<br>17.9 | 19.0;<br>18.7;<br>18.0 | 18.6;<br>18.9;<br>19.8 | 18.9;<br>18.7;<br>19.9 |

**Table 3:**
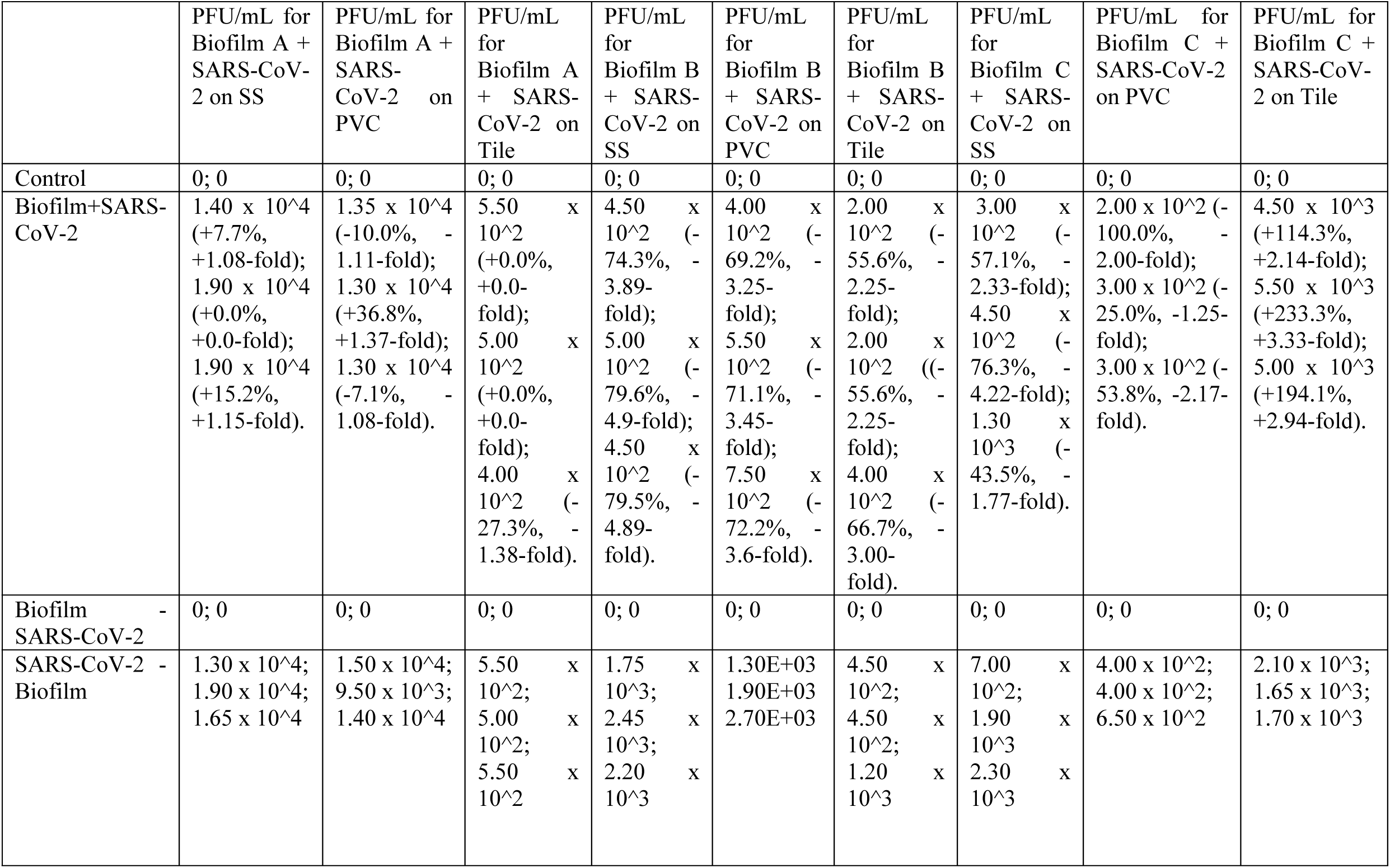
Data from the plaque assay analysis on the recovered SARS-CoV-2 Delta variant viral particles incubated with biofilm. Table 3 indicates PFU/mL numbers and the percentage and fold change from the initial inoculum (1.0 × 10^4^).

|  | PFU/mL for<br>Biofilm A +<br>SARS-CoV-2 on SS | PFU/mL for<br>Biofilm A +<br>SARS-CoV-2 on<br>PVC | PFU/mL<br>for<br>Biofilm A<br>+ SARS-CoV-2 on<br>Tile | PFU/mL<br>for<br>Biofilm B<br>+ SARS-CoV-2 on<br>SS | PFU/mL<br>for<br>Biofilm B<br>+ SARS-CoV-2 on<br>PVC | PFU/mL<br>for<br>Biofilm B<br>+ SARS-CoV-2 on<br>Tile | PFU/mL<br>for<br>Biofilm C<br>+ SARS-CoV-2 on<br>SS | PFU/mL for<br>Biofilm C +<br>SARS-CoV-2<br>on PVC | PFU/mL for<br>Biofilm C +<br>SARS-CoV-2 on Tile |
| --- | --- | --- | --- | --- | --- | --- | --- | --- | --- |
| Control | 0; 0 | 0; 0 | 0; 0 | 0; 0 | 0; 0 | 0; 0 | 0; 0 | 0; 0 | 0; 0 |
| Biofilm+SARS-<br>CoV-2 | 1.40 x 10<sup>4</sup><br>(+7.7%,<br>+1.08-fold);<br>1.90 x 10<sup>4</sup><br>(+0.0%,<br>+0.0-fold);<br>1.90 x 10<sup>4</sup><br>(+15.2%,<br>+1.15-fold). | 1.35 x 10<sup>4</sup><br>(-10.0%, -<br>1.11-fold);<br>1.30 x 10<sup>4</sup><br>(+36.8%,<br>+1.37-fold);<br>1.30 x 10<sup>4</sup><br>(-7.1%, -<br>1.08-fold). | 5.50 x<br>10<sup>2</sup><br>(+0.0%,<br>+0.0-<br>fold);<br>5.00 x<br>10<sup>2</sup><br>(+0.0%,<br>+0.0-<br>fold);<br>4.00 x<br>10<sup>2</sup><br>(-<br>27.3%, -<br>1.38-fold). | 4.50 x<br>10<sup>2</sup><br>(-<br>74.3%, -<br>3.89-fold);<br>5.00 x<br>10<sup>2</sup><br>(-<br>79.6%, -<br>4.9-fold);<br>4.50 x<br>10<sup>2</sup><br>(-<br>79.5%, -<br>4.89-fold). | 4.00 x<br>10<sup>2</sup><br>(-<br>69.2%, -<br>3.25-fold);<br>5.50 x<br>10<sup>2</sup><br>(-<br>71.1%, -<br>3.45-fold);<br>7.50 x<br>10<sup>2</sup><br>(-<br>72.2%, -<br>3.6-fold). | 2.00 x<br>10<sup>2</sup><br>(-<br>55.6%, -<br>2.25-fold);<br>2.00 x<br>10<sup>2</sup><br>(-<br>55.6%, -<br>2.25-fold);<br>4.00 x<br>10<sup>2</sup><br>(-<br>66.7%, -<br>3.00-fold). | 3.00 x<br>10<sup>2</sup><br>(-<br>57.1%, -<br>2.33-fold);<br>4.50 x<br>10<sup>2</sup><br>(-<br>76.3%, -<br>4.22-fold);<br>1.30 x<br>10<sup>3</sup><br>(-<br>43.5%, -<br>1.77-fold). | 2.00 x 10<sup>2</sup> (-<br>100.0%, -<br>2.00-fold);<br>3.00 x 10<sup>2</sup> (-<br>25.0%, -1.25-fold);<br>3.00 x 10<sup>2</sup> (-<br>53.8%, -2.17-fold). | 4.50 x 10<sup>3</sup><br>(+114.3%,<br>+2.14-fold);<br>5.50 x 10<sup>3</sup><br>(+233.3%,<br>+3.33-fold);<br>5.00 x 10<sup>3</sup><br>(+194.1%,<br>+2.94-fold). |
| Biofilm -<br>SARS-CoV-2 | 0; 0 | 0; 0 | 0; 0 | 0; 0 | 0; 0 | 0; 0 | 0; 0 | 0; 0 | 0; 0 |
| SARS-CoV-2 -<br>Biofilm | 1.30 x 10<sup>4</sup>;<br>1.90 x 10<sup>4</sup>;<br>1.65 x 10<sup>4</sup> | 1.50 x 10<sup>4</sup>;<br>9.50 x 10<sup>3</sup>;<br>1.40 x 10<sup>4</sup> | 5.50 x<br>10<sup>2</sup>;<br>5.00 x<br>10<sup>2</sup>;<br>5.50 x<br>10<sup>2</sup> | 1.75 x<br>10<sup>3</sup>;<br>2.45 x<br>10<sup>3</sup>;<br>2.20 x<br>10<sup>3</sup> | 1.30E+03<br>1.90E+03<br>2.70E+03 | 4.50 x<br>10<sup>2</sup>;<br>4.50 x<br>10<sup>2</sup>;<br>1.20 x<br>10<sup>3</sup> | 7.00 x<br>10<sup>2</sup>;<br>1.90 x<br>10<sup>3</sup><br>2.30 x<br>10<sup>3</sup> | 4.00 x 10<sup>2</sup>;<br>4.00 x 10<sup>2</sup>;<br>6.50 x 10<sup>2</sup> | 2.10 x 10<sup>3</sup>;<br>1.65 x 10<sup>3</sup>;<br>1.70 x 10<sup>3</sup> |

## Acknowledgements

The authors would like to thank USDA-NIFA 2020-67015-32330 grant for support of this study. We would like to thank Drs. Mick Bosilivac and Rong Wang (USDA-ARS-UAMRC, Clay Center-Nebraska) for providing meat packaging plant biofilms, and Dr. Ben Neuman (Texas A&M) for providing the original stocks of SARS-CoV-2 Delta variant for our study.

